# Multi-Parametric Phase-Cycled Balanced Steady-State Free Precession Brain Tissue Characterization in Relapsing-Remitting Multiple Sclerosis

**DOI:** 10.64898/2026.06.17.732900

**Authors:** Florian Birk, Benjamin Bender, Hamzeh Tesh, Anagha Deshmane, Tobias Lindig, Ulrike Ernemann, Klaus Scheffler, Rahel Heule

## Abstract

Quantitative MRI enables the detection of subtle microstructural alterations in normal-appearing white matter (NAWM) associated with pathological conditions such as multiple sclerosis. Quantitative metrics including R_1_, R_2_, and the bSSFP asymmetry index (AI) were evaluated in the WM of 20 relapsing-remitting multiple sclerosis (RRMS) patients and 10 healthy controls (HC). A multi-parametric framework based on a phase-cycled balanced steady-state free precession (pc-bSSFP) sequence was used. Diffusion tensor imaging–derived measures, including fiber-to-field angle, number of fiber orientations, and fractional anisotropy, were incorporated to assess parameter anisotropy. Statistical analysis was performed using linear mixed-effects models to test for group, ROI, and group-by-ROI effects for each metric, with ROI-specific group comparisons derived from the model. Significant main effects of group, ROI, and group-by-ROI interaction were observed for both R_1_ and R_2_, whereas for AI only the ROI effect reached significance (group p= 0.462; group-by-ROI p = 0.786). Fourteen of sixteen ROIs demonstrated significantly lower R_1_ and R_2_ values in RRMS compared with HC. No ROI showed significant differences in AI. In conclusion, pc-bSSFP–based relaxometry reveals predominantly white matter alterations in RRMS, while enabling a comprehensive whole-brain assessment that also encompasses gray matter.

## Introduction

Multiple sclerosis (MS) is one of the most prevalent inflammatory and neurodegenerative diseases of the central nervous system (CNS) with multifactorial causes underlying its development and progression. The myelinated axons in the CNS, which form the core architecture for fast signal transmission in white matter (WM), may be targeted by activated immune cells associated with widespread microglial activation or may experience reduced support from oligodendroglial cells [1, 2]. While the detection of focal lesions is a clear indicator of MS, it is not necessarily specific to clinical disability measures. Consequently, there has been increased interest in the characterization of non-lesional tissue alterations. Diffusely abnormal white matter (DAWM) is characterized by subtle non-focal increases in signal intensity on proton density and *T*_2_-weighted images, with values intermediate between normal-appearing WM (NAWM) and focal lesions, spatially occurring either adjacent to or distant from lesions, and has been reported across all clinical phenotypes of MS [1]. Moreover, DAWM has been observed already in early disease onset [3], with prevalences of approximately 25% in patients with clinically isolated syndromes [4] and up to 73% in relapsing-remitting MS (RRMS) [5]. Thus, disease progression independent of relapse activity has been identified as a key contributor to disability accumulation [6, 7].

Quantitative magnetic resonance imaging (qMRI) aims to provide objective measures of biophysical tissue properties that reflect subtle changes in pathological tissue. In particular, longitudinal (*T*_1_) and transverse (*T*_2_) relaxation times, magnetization transfer, as well as diffusion tensor imaging (DTI) metrics such as fractional anisotropy (FA) have been reported to correlate with ex vivo histopathology [8] and to reflect the severity of pathological changes in vivo in chronic MS phenotypes [9]. Increased *T*_1_ and *T*_2_ values and decreased FA values in DAWM are associated with extensive axonal loss and reduced myelin density in patients with chronic MS [8] and multi-shell diffusion imaging has further revealed changes in myelin and axonal integrity in the NAWM of MS patients [10]. Moreover, *T*_2_ relaxation values in DAWM were reported to be intermediate between NAWM and *T*_1_w-hypointense focal lesions [5], and *T*_1_ relaxation times were shown to be sensitive to microstructural changes in early RRMS [11].

Quantitative MRI faces challenges due to the need to acquire multiple weighted MR contrasts, leading to long scan times, and is often limited to measuring a single biophysical parameter. Recent advances have focused on the development of multi-parametric (MP) methods, which provide multiple co-registered quantitative maps within clinically relevant scan times. Dif-ferent frameworks have been suggested for rapid multi-parametric 3D whole-brain tissue characterization with focus on relaxometry. These include steady-state approaches, e.g. multi-parameter mapping (MPM) [12] or phase-cycled balanced SSFP (pc-bSSFP) [13, 14, 15], and transient-state imaging, e.g. magnetic resonance fingerprinting (MRF) [16], MR multitasking [17], or magnetic resonance spin tomography in the time domain (MR-STAT) [18]. Recent studies have demonstrated that MP-mapping may be used to monitor subtle changes within the NAWM and plaque dynamics, reflecting tissue repair or disease progression [19], and may provide complementary tissue information through multiple estimated parameters, improving the differentiation between patients with MS and healthy controls (HC) [20, 21]. In addition, traditional DTI allows tract-specific analysis tailored to important WM bundles affected by RRMS [22, 23]. Phase-cycled bSSFP imaging offers high SNR efficiency and strong sensitivity to the intra-voxel frequency content reflecting tissue microstructure [24, 25]. While characteristic banding artifacts of bSSFP contrasts may hinder direct anatomical interpretation, phase-cycling enables effective sampling of the bSSFP frequency response profile, opening up versatile applications for relaxometry [13, 14, 15, 26, 27], fat fraction mapping [28], quantification of diffusion metrics [29], or direct mapping of the profile asymmetries using the asymmetry index (AI) related to intravoxel frequency distributions [25].

To the best of our knowledge, this study is the first to demonstrate the application of pc-bSSFP whole-brain imaging for fast multi-parametric relaxometry mapping in patients with RRMS in comparison to HC. Investigating the repeatability of the proposed method and the WM fiber orientation dependence of the derived quantitative parameters in combination with tractography-derived WM bundle analysis in MS-relevant structures provides a comprehensive overview of the potential use of multi-parametric pc-bSSFP frameworks to assess tissue integrity in neurodegenerative diseases, such as MS.

## Methods

### Subjects

MRI experiments were conducted using a clinical MR scanner with a field strength of 3 T (Magnetom Vida, Siemens Healthineers, Erlangen, Germany) and a manufacturer-built 20-channel receive head array coil. All experiments were performed in accordance with local ethical guidelines and written informed consent was obtained from all participants prior to the MR measurement. Ten HC (age range/mean/standard deviation 27–47/33.4/6.9 years) and 20 age-matched RRMS subjects (age range/mean/standard deviation: 26–47 years/34.5/5.9 years) were included in this study. All RRMS subjects were measured as part of a clinical routine MR examination. To assess the repeatabil-ity of the proposed methods, five HC subjects underwent baseline and direct follow-up measurements at the same MR scanner using an identical MR scan protocol. The follow-up measurements were further subdivided into intra-session measurements performed in three HC subjects where the entire protocol was repeated a second time (repeat) and inter-session measurements performed in two different HC subjects where the subject was taken out of the MR scanner and repositioned for the next scan (reposition).

### Data Acquisition

The acquisition protocol included a *T*_1_-weighted magnetization-prepared rapid gradient-echo (MPRAGE) sequence [30] and a *T*_2_-weighted fluid-attenuated inversion recovery (FLAIR) [31] sequence for brain tissue segmentation and automatic lesion segmentation in RRMS patients, respectively. Quantitative MRI sequences included a 4D whole-brain phase-cycled bSSFP measurement with 12 linear phase increments (*ϕ*) uniformly distributed in the range of 0–2 *π* for multi-parametric mapping, a vendor transmit field 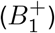 mapping [32] sequence to correct for 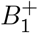 inhomogeneities, and a single-shell DTI sequence with 30 diffusion directions (b-values = 0, 1000 s mm^−2^). Detailed protocol parameters can be found in Table 1.

**Table 1.** Study MR protocol.

| Sequence | Anatomical $T_1$<br>3D T1-weighted<br>MPRAGE | Anatomical $T_2$<br>3D T2-weighted<br>FLAIR | PC-bSSFP<br>4D pc-bSSFP | $B_1^+$<br>2D multi-slice<br>TurboFLASH<br>with/without pre-<br>conditioning RF<br>pulse | DTI<br>2D multi-slice<br>single-shot SE-EPI |
| --- | --- | --- | --- | --- | --- |
| FOV | 256 × 256 × 176 mm<sup>3</sup> | 256 × 256 × 176 mm<sup>3</sup> | 256 × 256 × 176 mm<sup>3</sup> | 256 × 256 × 176 mm<sup>3</sup> | 256 × 256 × 176 mm<sup>3</sup> |
| Resolution | 1 × 1 × 1 mm<sup>3</sup> | 0.9 × 0.9 × 0.9 mm<sup>3</sup> | 1.3 × 1.3 × 1.3 mm<sup>3</sup> | 2.4 × 2.4 × 3 mm<sup>3</sup> | 2 × 2 × 2 mm<sup>3</sup> |
| # of slices | 176 | 192 | 128 | 30 | 80 |
| TR | 2000 ms | 5000 ms | 4.8 ms | 14.2 s | 5800 ms |
| TE | 3.26 ms | 1800 ms | 2.4 ms | 2.4 ms | 73 ms |
| TI |  | 386 ms |  |  |  |
| Flip angle | 8° | variable | 15° | 8° | 90° excitation; 180°<br>refocusing |
| Diffusion<br>direc-<br>tions/Number<br>of phase cy-<br>cles | | | 12 phase cycles;<br>256 dummy prepa-<br>ration pulses | | Monopolar; 30<br>diffusion directions<br>($b = 1000 \text{ s mm}^{-2}$);<br>two runs<br>($b = 0 \text{ s mm}^{-2}$)<br>with opposite<br>phase encoding<br>directions |
| Parallel Imag-<br>ing | GRAPPA accel-<br>eration 2 (in-<br>plane/through-<br>plane: 2/1) | GRAPPA accel-<br>eration 2 (in-<br>plane/through-<br>plane: 2/1) | GRAPPA accel-<br>eration 2 (in-<br>plane/through-<br>plane: 2/1) |  | Simultaneous<br>multi-slice (SMS)<br>acceleration 2 |
| Scan time | 4 min and 40 s | 4 min and 57 s | 10 min and 13 s | 29 s | 3 min and 30 s |

### Data Processing

The processing of the acquired data can be divided into three main parts, as shown in Figure 1a.

**Figure 1.**
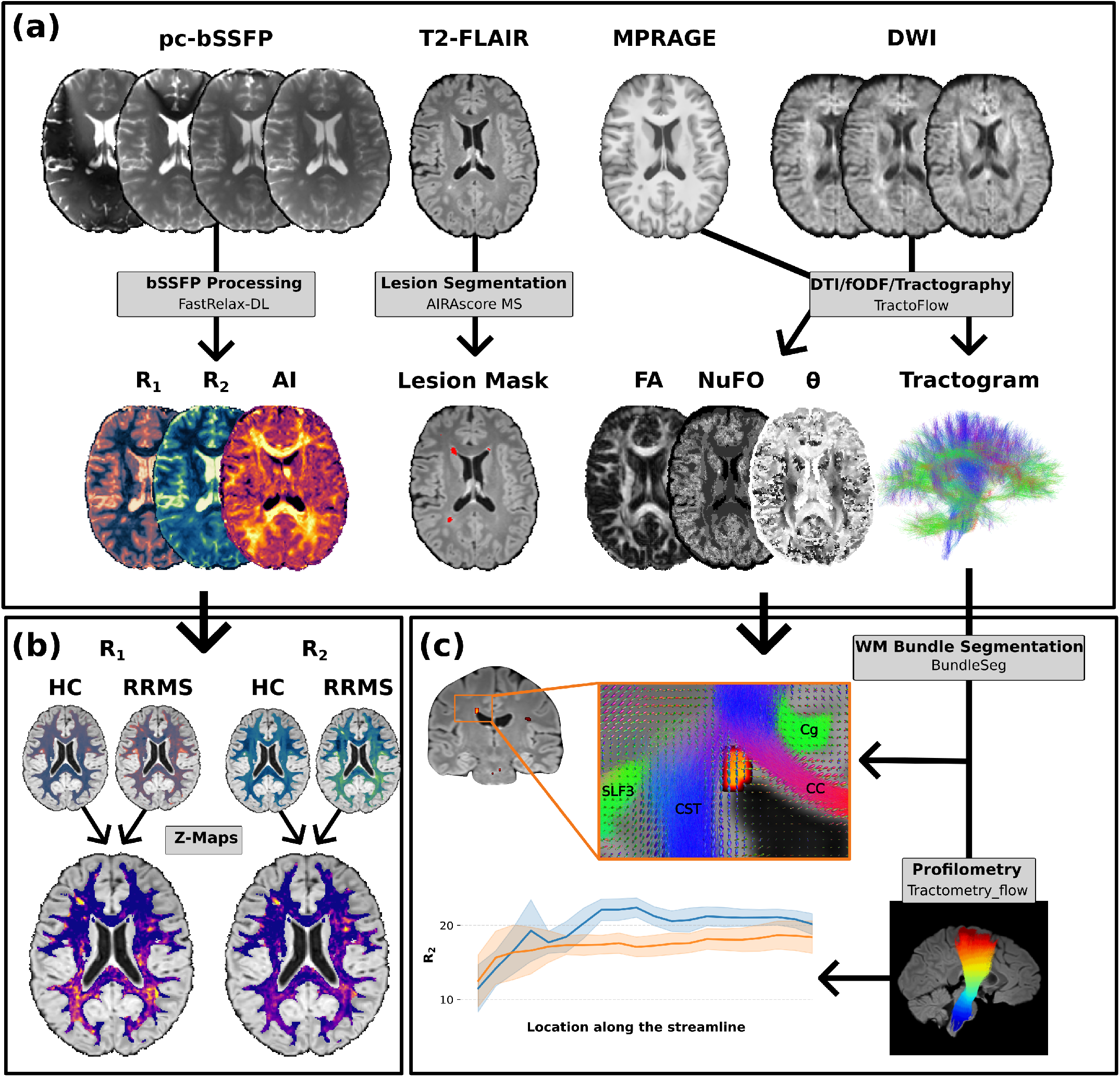
Data processing scheme and tools employed in this work. (a) The 4D pc-bSSFP data was used to simultaneously estimate both *R*_1_ and *R*_2_ relaxation maps using the FastRelax-DL MP-framework [27] and to extract the bSSFP-specific asymmetry index (AI). Lesions in the *T*_2_ -FLAIR images were automatically segmented using *AIRAscore* (version 2.4.0, AIRAmed GmbH, Tübingen) [33]. Anatomical *T*_1_ -weighted MPRAGE contrasts and DWI data were used as inputs to the *TractoFlow* pipeline [34], yielding FA, NuFO, and fiber-to-field angle *θ* derived from the primary peak of the fiber orientation distribution function, and a whole-brain WM tractogram as outputs. (b) To further assess voxelwise differences between HC subjects and RRMS patients, relaxation rate maps were registered into MNI space for z-score estimation in WM. (c) Whole-brain tractograms were further segmented into WM bundles using the tool *BundleSeg* [35]. Profilometry along specific fibers was achieved using the *tractometry_flow* pipeline [36].

#### Tractography using TractFlow

Single-shell diffusion weighted imaging (DWI) and anatomical *T*_1_-weighted MPRAGE data were used as input to the *TractoFlow* pipeline [34]. DWI preprocessing steps included initial denoising, followed by correction of magnetic susceptibility- and motion-induced artifacts with slice-wise outlier detection, and subsequent brain extraction based on distortion-free images. Subsequent DWI pre-processing steps consisted of N4 bias field correction, image cropping, normalization, and final resampling of the DWI images to 1 mm isotropic resolution to match the spatial resolution of the *T*_1_-weighted anatomical image. The *TractoFlow* pipeline fitted the DWI data to the DTI tensor model to calculate the FA. To account for crossing fibers, the fiber orientation distribution function (fODF) was computed by first estimating the fiber response function (FRF) from voxels with FA *>* 0.7. Constrained spherical deconvolution was then applied to the single-shell DWI data using the FRF as kernel to derive the fODF in each voxel. Based on the fODF peaks, the number of fiber orientations (NuFO) within a voxel and the fiber-to-field angle referring to the primary peak direction of the fODF were extracted. Preprocessing of the *T*_1_-weighted data included denoising, N4 bias field correction, brain extraction, cropping, and registration to the DWI images. The preprocessed *T*_1_-weighted data were segmented into WM, gray matter (GM), and cerebrospinal fluid (CSF). The resulting masks were used as tracking maps for tractography, with the WM segmentation serving as the seeding region. The constrained particle filter tracking algorithm was used to generate streamlines and whole-brain tractograms. As shown in Figure 1c, the whole-brain tractogram was further segmented into WM pathways using the *BundleSeg* package [35] and the ICBM152 nonlinear asymmetric 2009a atlas [37] as reference.

#### Relaxometry and Asymmetry Index

Preprocessing of the complex-valued 4D phase-cycled bSSFP data included intra-registration of the acquired 12 3D phase-cycles to a middle phase cycle in the bSSFP passband (*ϕ* = 165^*°*^), phase correction to remove receiver-related offsets, followed by ringing artifact removal [38]. The three lowest-order SSFP configurations (*F*_1_, *F*_0_, and *F*_1_) were obtained based on a discrete 12-point Fourier transform of the complex-valued pc-bSSFP data. The *F*_1_, *F*_0_, and *F*_1_ configurations and the co-registered median-filtered 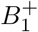 map served as input to a deep learning-based relaxometry framework for combined *R*_1_(= 1*/T*_1_) and *R*_2_(= 1*/T*_2_) relaxation rate estimation [27]. The AI was calculated based on the normalized *B*_0_-corrected bSSFP frequency response as AI = (*h*_*p*_ *− h*_*n*_)*/*(*h*_*p*_ + *h*_*n*_), where *h*_*p*_ and *h*_*n*_ represent the signal peaks at positive and negative frequency offsets [25]. The mean pc-bSSFP magnitude was then registered to the final anatomical *T*_1_-weighted reference of the *TractoFlow* pipeline using ANTs and the transformation was applied to all quantitative metrics (*R*_1_, *R*_2_, AI).

#### Automatic Lesion Segmentation

For RRMS data, high-resolution *T*_2_-FLAIR images were resampled to 1 mm isotropic resolution and processed for focal lesion segmentation using AIRAscore (version 2.4.0, AIRAmed GmbH, Tübingen) [33]. Segmentation was performed within a research pipeline using NIfTI inputs and outputs, rather than the vendor’s clinical workflow, to enable downstream quantitative analyses.

#### MNI Space

The obtained quantitative pc-bSSFP metrics (*R*_1_, *R*_2_, AI) and the *T*_2_-FLAIR data with lesion segmentations of all subjects were registered to a common reference space. The *T*_1_-weighted data derived from the *TractoFlow* output was registered to the ICBM152 atlas space using *SynthMorph* [39]. The regularization parameter *λ* was set to 0.25 (*λ*∈ [0, 1]; 0 indicating maximal deformation, and 1 indicating no deformation). Subsequently, transformations from the *T*_2_-FLAIR or bSSFP space to the *TractoFlow T*_1_-weighted space, along with its *SynthMorph* displacement field to the ICBM152 *T*_1_-weighted space, were combined and applied using ANTs. Linear interpolation was used for all anatomical and quantitative data, while generic label interpolation was used for the lesion segmentations. An HC atlas was generated in the common space for both *R*_1_ and *R*_2_ relaxation rates. The mean (HC_*µ*_) and standard deviation (HC_*σ*_) of all HC baseline measurements were computed, and voxelwise z-scores for RRMS subjects were calculated as z-score = (RRMS *−* HC_*µ*_)*/*HC_*σ*_.

### Data Analysis

After data quality control and age-matching to the HC, 16 RRMS subjects were included into the final data analysis (age range/mean/std: 26–47/34.4/6.3 years). If not stated otherwise, the analysis was performed in the subject space and pooled across HC and RRMS cohorts.

The scan-rescan repeatability of the proposed MP framework was assessed using HC baseline and follow-up measurements acquired in five subjects. For each metric (*R*_1_, *R*_2_, AI), mean values per measurement (baseline or follow-up) and subject were calculated in WM and GM regions of interest (ROIs), i.e. corpus callosum (CC), cerebellar WM, frontal WM, temporoparietal WM, occipital-parietal WM, cerebral WM, cerebral GM, thalamus, and caudate, and utilized for Bland–Altman analyses.

In this study, NAWM is used to denote both normal-appearing and diffusely abnormal WM, as no distinction between these tissue classes was made. The orientation dependence in NAWM, excluding voxels from the lesion masks in RRMS subjects, was assessed in baseline HC and age-matched RRMS cohorts. A WM mask was derived from the anatomical *T*_1_-weighted data by including voxels with a WM probability *>* 90%. WM vox-els were further binned by fiber-to-field angle *θ* ranging from 0–90° in 7.5° increments with additional thresholding applied to FA (0.5, 1.0], *R*_1_ [1.0, 3.0] 1/s, and *R*_2_ [13.0, 29.0] 1/s. The mean and standard deviation (SD) across both cohorts were calculated for each metric and bin. The strength of the orientation dependence, i.e. the parameter anisotropy, was calculated as ((R_i,max_ *−* R_i,min_)*/*(R_i,max_ + R_i,min_)) *·* 100 (with i = 1, 2 for *R*_1_ and *R*_2_) and (AI_max_ *−* AI_min_) *·* 100 for the inherently normalized AI metric.

To analyze group-wise differences along specific WM bundles, all metrics of interest were combined with the *BundleSeg* WM tracts as input for the tractometry flow pipeline [36], which segmented the WM bundles of each subject into 20 equidistant segments (see Figure 1c) and computed the mean values of the quantitative metrics as well as the lesional volume for each of these segments. The following WM bundles were included in the tract profile analysis: superior longitudinal fasciculus (SLF) subdivided into three compartments (SLF1, SLF2, SLF3), inferior longitudinal fasciculus (ILF), inferior fronto-occipital fasciculus (IFOF), corticospinal tract (CST), and arcuate fasciculus (AF).

### Statistical Analysis

Quantitative MRI metrics (*R*_1_, *R*_2_, AI) were compared between the age-matched HC and RRMS participants across all 16 ROIs using linear mixed-effects models (LMM) with one LMM fit per metric. For each metric, the primary model was value ∼ group *×* ROI + age + gender + (1 | subject), with group, ROI, age, and gender as fixed effects and a subject-specific random intercept. The main inferential focus was the group-by-ROI interaction, which tests whether HC-RRMS differences vary by region rather than remaining constant across the brain. ROI-specific HC versus RRMS two-tailed uncorrected p values (p_uncorr_) were obtained from linear contrasts combining the main group effect and the ROI-specific interaction term. Effect sizes were summarized using Cohen’s d. Anisotropy differences between HC and RRMS were assessed at each NuFO level using two-sided Welch t tests assuming unequal variances. For both analyses, multiple testing was controlled using the Benjamini–Hochberg false discovery rate (FDR) procedure, and FDR-corrected p values were reported as p_corr_ with significance set at p_corr_ *<* 0.05.

## Results

### Repeatability and Orientation Dependence

The fiber-to-field angle dependency of *R*_1_, *R*_2_, and AI is illustrated in Figure 2a-c representatively for two individual HC subjects undergoing intra-session (repeat, blue curves) and inter-session (reposition, green curves) as well as for the HC cohort mean across all baseline measurements (black curves). High anisotropy repeatability is observed for *R*_2_ and AI whereas *R*_1_ exhibits somewhat higher variability due to its lower degree of anisotropy. The ROI-based Bland-Altman analysis of the scan-rescan repeatability in HC subjects yielded mean biases of 1.48% (SD = 3.43%), 2.62% (SD = 3.59%), and 0.21% (SD = 0.56%) for the assessed ROIs in *R*_1_, *R*_2_, and AI, respectively (cf. Figure 2d-e).

**Figure 2.**
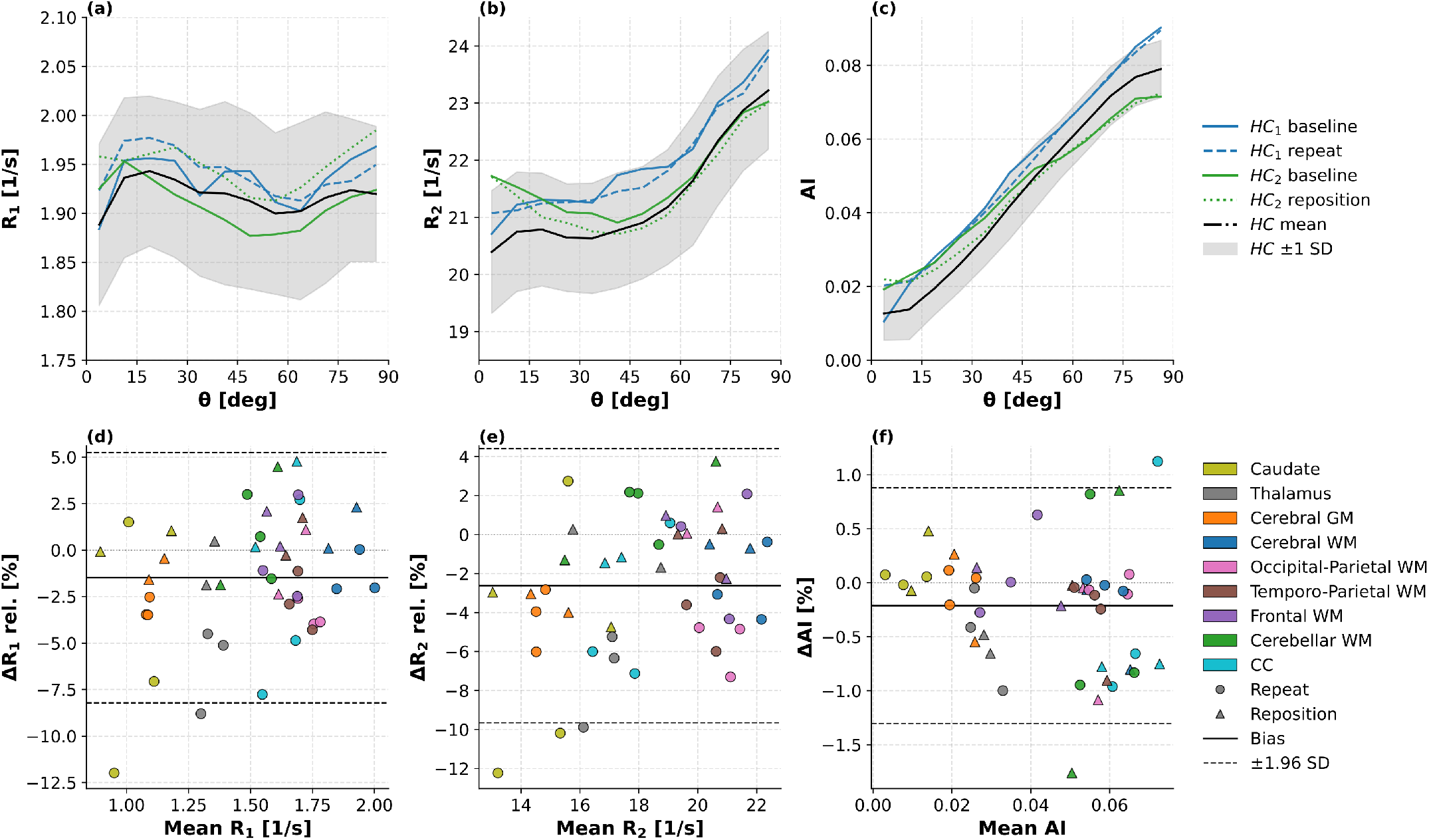
Repeatability analysis of the relaxation rates *R*_1_ and *R*_2_, and the asymmetry metric AI in HC. (a)-(c) Mean anisotropy values of *R*_1_ (a), *R*_2_ (b), and AI (c) versus fiber-to-field angle *θ* in WM pooled across all ten baseline HC assessments (black curves, ± SD in gray) as well as in two individual HC subjects undergoing intra-session (repeat, blue curves) and inter-session (reposition, green curves) repeatability analysis are plotted. Data are binned by *θ* using a step size of 7.5°and thresholded by FA (0.5, 1.0]. (d)-(f) Bland-Altman plots including GM (caudate, thalamus, cerebral), and WM (cerebral, occipital-parietal, temporoparietal, frontal, cerebellar, CC) ROIs are shown for *R*_1_ (d), *R*_2_ (e), and AI (f). The relative difference in percentage was derived for each parameter ROI as ((follow-up *−* baseline)*/*mean) *·* 100 with mean = (follow-up *−* baseline)*/*2 for *R*_1_ and *R*_2_, and 1 for AI. The mean relative difference (bias) and the 95% limits (mean bias ± 1.96 SD) are plotted in solid and dashed black horizontal lines, respectively. HC subjects with repeated and reposition measurements are shown as circles and triangles, respectively.

Comparing the mean NAWM anisotropy of the quantitative metrics between HC (blue curves) and RRMS subjects (orange curves) in Figure 3, three main observations can be made: (1) mean NAWM *R*_1_ and *R*_2_ values in RRMS are lower than in HC across all *θ* bins, (2) the strong orientation dependence of *R*_2_ and AI decreases with a higher number of intra-voxel fiber orientations (left: NuFO = 1, middle: NuFO = 2, right: NuFO = 3), and (3) the *R*_2_ and AI anisotropy appears reduced for the RRMS cohort compared to the HC cohort. The quantitative assessment of the anisotropy strength resulted in values decreasing from 5.96% (SD = 1.64%) and 5.45% (SD = 0.76%) at NuFO = 1 to 2.85% (SD = 0.97%) and 1.74% (SD = 0.47%) at NuFO = 3 for *R*_2_ and AI, respectively, in case of HC. For RRMS, the calculated anisotropy decreased from 4.94% (SD = 1.64%) and 4.86% (SD = 0.75 %) at NuFO = 1 to 2.03% (SD = 0.75%) and 1.50% (SD = 0.33%) at NuFO = 3 for *R*_2_ and AI, respectively.

**Figure 3.**
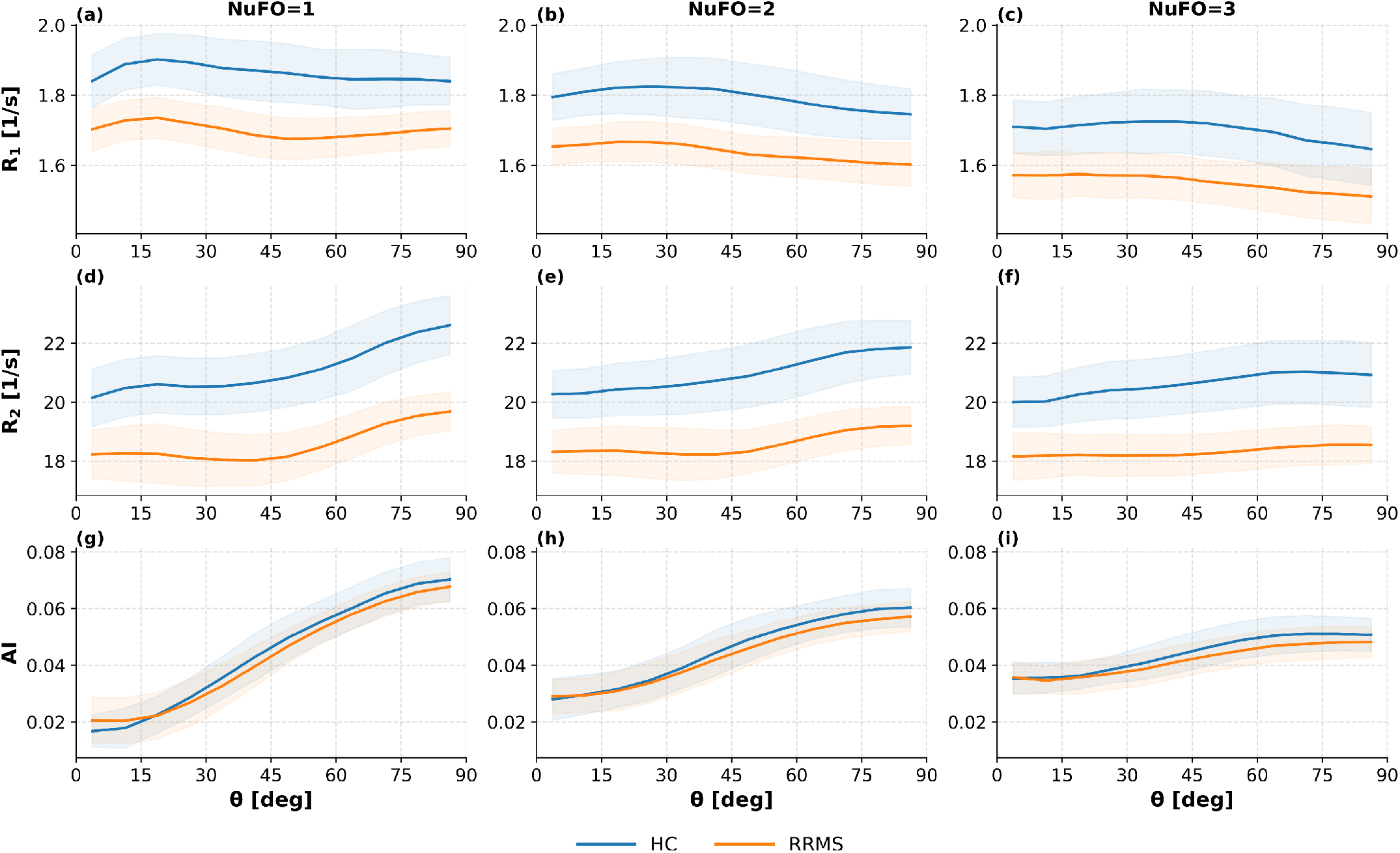
Parameter anisotropy dependence on the number of fiber orientations and cohort. The mean (± SD) anisotropy of *R*_1_ (a-c), *R*_2_ (d-f), and AI (g-i) versus fiber-to-field angle *θ* in NAWM for age-matched HC and RRMS cohorts is shown in blue and orange, respectively. Anisotropy analysis is further subdivided into one (first column), two (second column), and three (third column) intra-voxel fiber orientations (NuFO).

### MNI Space Analysis

Figure 4 presents quantitative maps of WM *R*_1_ and *R*_2_ for three axial slices (columns 1–3) of a representative RRMS subject in comparison to the mean HC atlas in MNI space, revealing WM lesions and diffuse WM alterations. Focal lesions, indicated as red segments overlaid on the anatomical *T*_2_-FLAIR data in the last row, appear hyperintense in the quantitative maps. This is quantitatively supported by the calculated mean absolute z-score values of 3.61 (SD = 2.82) and 3.56 (SD = 2.31) for *R*_1_ and *R*_2_, respectively, in the whole-brain lesion mask of the presented RRMS subject. Moreover, lesion-like WM regions with z-score values exceeding 3, but appearing normal on the *T*_2_-FLAIR data, are visible, as well as WM regions with moderately increased z-scores in the range 2–3, suggesting diffuse WM change.

**Figure 4.**
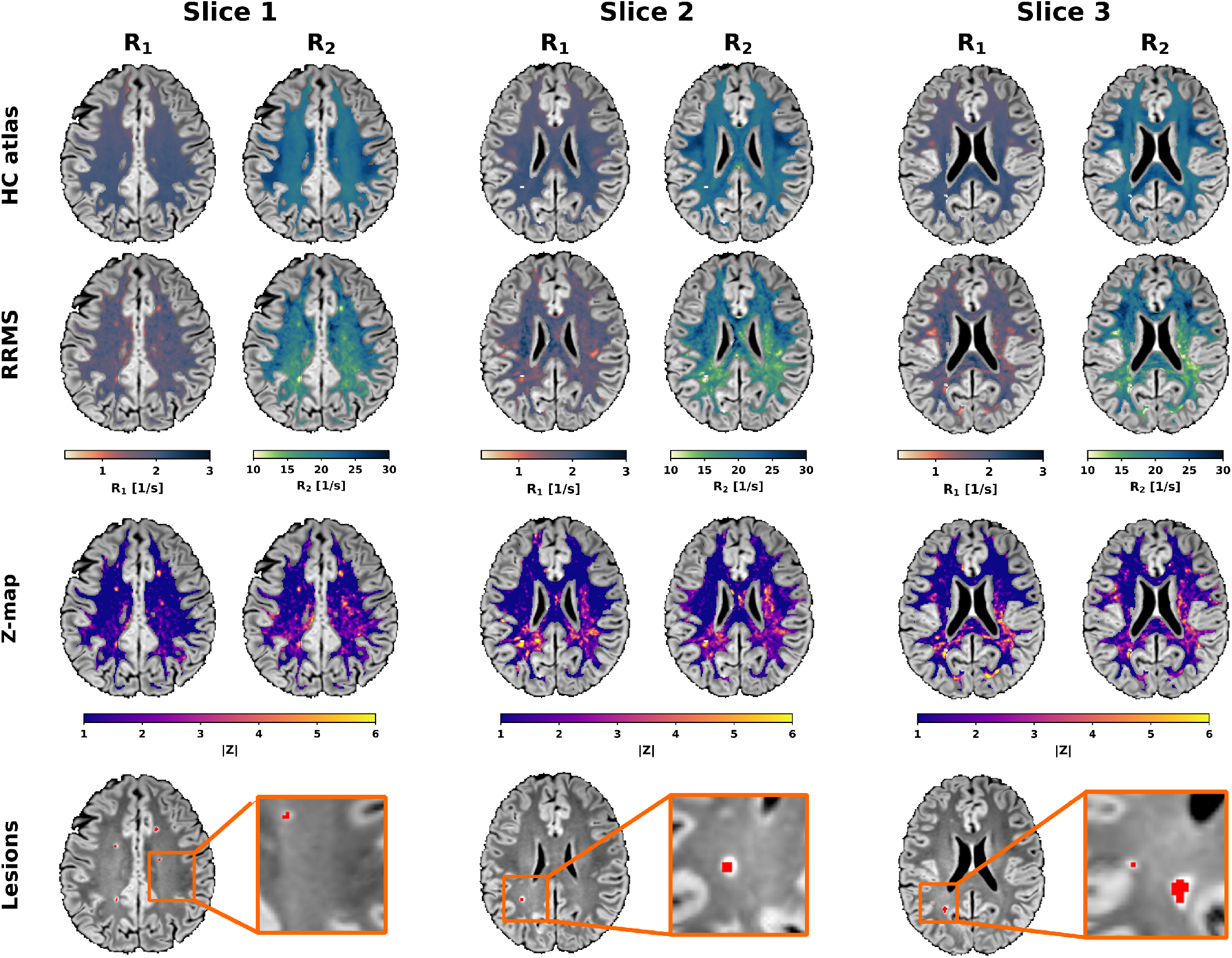
MNI space analysis of a representative RRMS patient (30 years, female). Three exemplary axial slices are shown for the HC atlas (HC_*µ*_) (first row) and for one RRMS subject (second row) with WM-segmented *R*_1_ and *R*_2_ maps overlaid onto the anatomical *T*_2_ -FLAIR image of the RRMS subject. The absolute z-score maps in the third row are calculated as |Z| = |(RRMS *−* HC_*µ*_)*/*HC_*σ*_| to emphasize the magnitude of voxelwise deviation from the normative atlas (abnormality burden) independent of direction. The last row displays the lesion masks transformed into MNI space and overlaid in red on the *T*_2_ -FLAIR, along with zoomed-in views of lesion-proximal regions.

### Profilometry

Figure 5 illustrates tract-specific analysis of the agematched HC and RRMS cohorts for two representative WM tracts, CST (orange, red) and SLF1 (cyan, blue), in the left and right hemispheres. Streamline profiles of NuFO, FA, and fiber-to-field angle *θ* show a high degree of agreement between mean HC and RRMS values, suggesting minimal confounding effects of tissue anisotropy. An increase in FA is associated with reduced NuFO values along the streamlines. For both left and right CST bundles, mean *R*_1_ and *R*_2_ values are reduced in RRMS relative to HC with largest differences of -0.3 1/s and -5.3 1/s for *R*_1_ and *R*_2_ in the mid-segments 10–12, respectively. This divergence co-incides with higher FA values in these segments. In the left and right SLF1, differences in mean *R*_1_ and *R*_2_ values between RRMS and HC are most pronounced in the posterior segments and gradually converge toward the anterior direction, with maximum differences of -0.34 1/s and -5.1 1/s in segments 1–5 relative to the mean HC profiles. The mean AI demonstrates similar values for HC and RRMS in CST but is reduced for both left and right SLF1 bundles in RRMS relative to HC along the streamlines.

**Figure 5.**
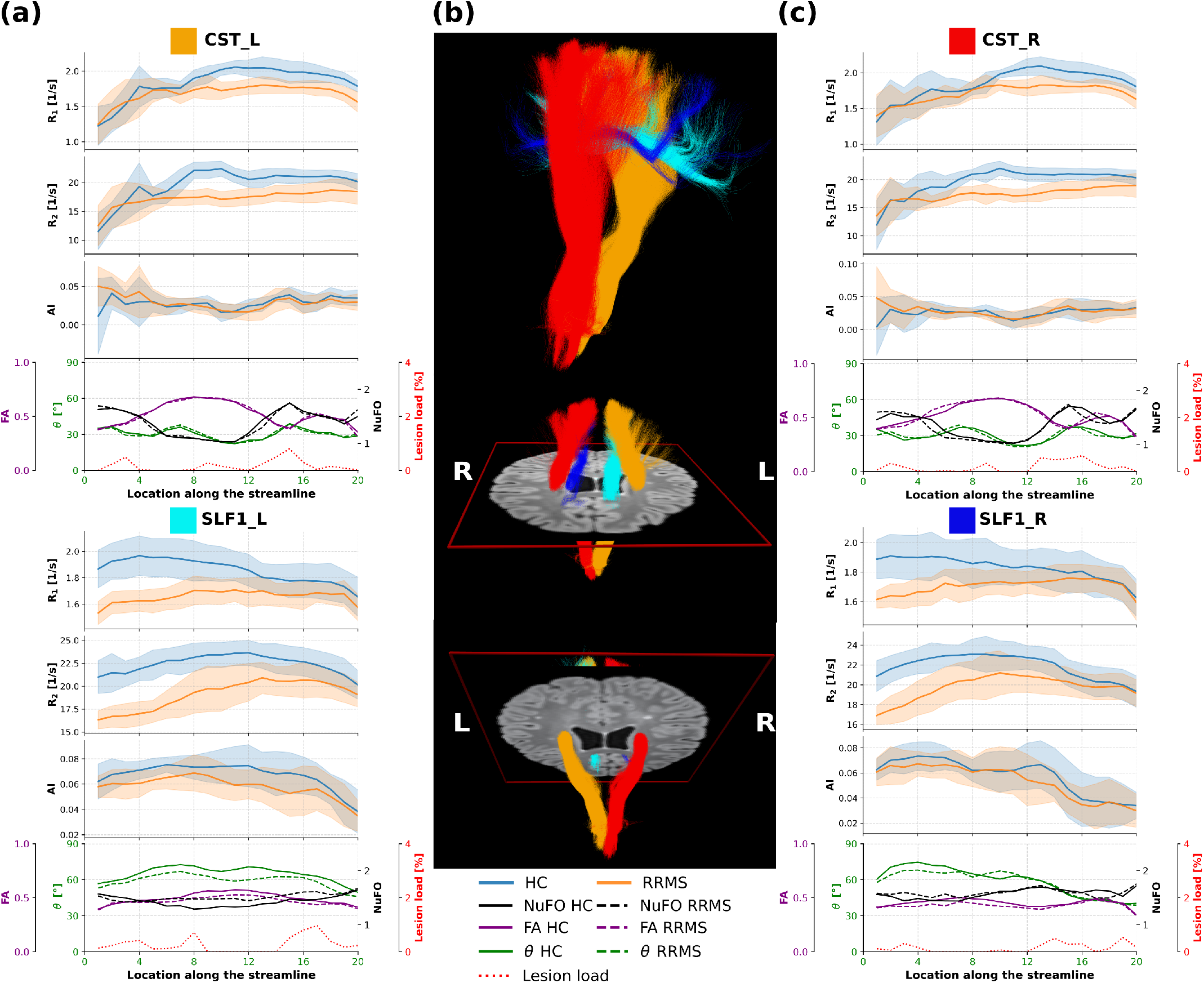
Multi-parametric tract-specific profilometry in age-matched HC and RRMS cohorts. In (a) and (c), tract-specific mean values of *R*_1_, *R*_2_, and AI, as well as of diffusion metrics, i.e. FA, fiber-to-field angle *θ*, and NuFO, and of the lesion load are plotted for the left (a) and right (c) hemispheres of the corticospinal tract (CST) and the superior longitudinal fasciculus (SLF1). The mean diffusion metrics assessed in HC and RRMS are displayed as solid and dashed curves, respectively. The middle column (b) presents a 3D rendering of the left and right CST (left = orange; right = red) and SLF1 (left = cyan; right = blue) WM tracts passing through an axial *T*_2_ -FLAIR image in a representative RRMS patient (female, 29 years).

### Statistical Analysis

As shown in Table 2, LMMs revealed significant group effects, ROI effects, and group-by-ROI interaction for *R*_1_ and *R*_2_. For AI, the group effect (p = 0.462) and group-by-ROI interaction (p = 0.786) were not significant. Age and gender effects were not significant across all metrics (age: p = 0.572 to 0.683; gender: p = 0.172 to 0.765). Table 3 and Figure 6 show significant *R*_1_ differences in all ROIs except cerebellar WM (p_uncorr_ = 0.200, p_corr_ = 0.200) and caudate (p_uncorr_ = 0.162, p_corr_ = 0.173) and significant *R*_2_ differences in all ROIs except frontal WM (p_uncorr_ = 0.107, p_corr_ = 0.114) and caudate (p_uncorr_ = 0.152, p_corr_ = 0.152). For AI, no ROI showed statistical significance before or after FDR correction. p_uncorr_ values ranged from 0.151 to 0.960 and p_corr_ values from 0.937 to 0.960. Cohen’s d indicated that the main cohort effect was negative for *R*_1_ and *R*_2_ in the ROIs that survived FDR correction, consistent with lower RRMS values than HC. The largest effect sizes were seen in temporo-parietal WM (d = -2.94), occipital-parietal WM (d = -2.83) and SLF3 (d = -2.52) for *R*_1_ and the ILF (d = - 4.25), temporo-parietal WM (d = -4.03), and thalamus (d = -3.57) for *R*_2_. AI showed smaller and less consistent effect sizes (d = -0.46 to 0.41), in line with the absence of significant ROI-level effects. For the anisotropy analysis, the AI p_uncorr_-values increased from NuFO = 1 to NuFO = 3 (p_uncorr_ = 0.067-0.173) and remained non-significant after FDR correction (p_corr_ = 0.108-0.173), whereas for *R*_2_ p_uncorr_-values decreased from NuFO = 1 to NuFO = 3 (p_uncorr_ = 0.138-0.037) but FDR correction increased the Nufo = 3 value to 0.055, so none met the significance threshold. Overall, mean RRMS NAWM values for *R*_1_, *R*_2_, and AI (1.65 1/s, 18.71 1/s, and 0.049, respectively) lay between the corresponding mean HC WM values (1.80 1/s, 21.28 1/s, 0.052) and the mean lesional values (1.18 1/s, 12.46 1/s, 0.043).

**Table 2.**
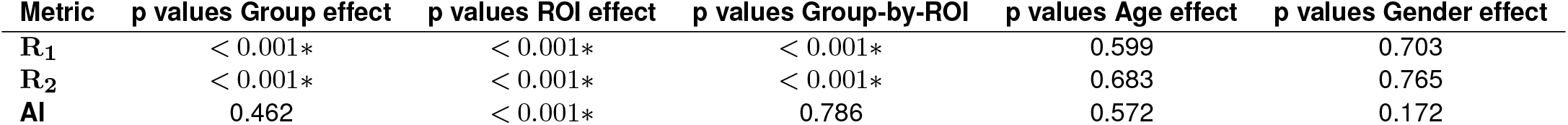
Primary linear mixed-effects model results for *R*_1_, *R*_2_, and AI, including fixed-effect p values for group, ROI, group-by-ROI, age, and gender. Ten HC and 16 age-matched RRMS with 16 ROIs per subject resulted in 416 observations per metric.

| Metric | p values Group effect | p values ROI effect | p values Group-by-ROI | p values Age effect | p values Gender effect |
| --- | --- | --- | --- | --- | --- |
| $R_1$ | < 0.001* | < 0.001* | < 0.001* | 0.599 | 0.703 |
| $R_2$ | < 0.001* | < 0.001* | < 0.001* | 0.683 | 0.765 |
| AI | 0.462 | < 0.001* | 0.786 | 0.572 | 0.172 |

**Table 3.** ROI-specific HC versus RRMS results for *R*_1_, *R*_2_, and AI in NAWM, including mean values, Cohen’s d, uncorrected (p_uncorr_) and corrected (p_corr_) p values. False discovery rate (FDR) was controlled using the Benjamini–Hochberg procedure. Statistical significance was defined as p_corr_ *<* 0.05 and is indicated by an asterisk (*).

| | Mean HC ± SD | | | Mean RRMS ± SD | | | Cohen's d | | | $p_{\text{uncorr}} (p_{\text{corr}})$ | | |
| --- | --- | --- | --- | --- | --- | --- | --- | --- | --- | --- | --- | --- |
| | $R_1$ | $R_2$ | AI | $R_1$ | $R_2$ | AI | $R_1$ | $R_2$ | AI | $R_1$ | $R_2$ | AI |
| **Caudate** | 1.22 ± 0.07 | 19.16 ± 0.84 | 0.01 ± 0.01 | 1.18 ± 0.05 | 18.58 ± 1.07 | 0.01 ± 0.01 | -0.76 | -0.58 | -0.21 | 0.162 (0.173) | 0.152 (0.152) | 0.776 (0.937) |
| **Thalamus** | 1.51 ± 0.03 | 19.93 ± 0.92 | 0.03 ± 0.01 | 1.39 ± 0.06 | 16.91 ± 0.80 | 0.03 ± 0.01 | -2.42 | -3.57 | -0.07 | <0.001 (<0.001*) | <0.001 (<0.001*) | 0.878 (0.937) |
| **Cerebral GM** | 1.27 ± 0.04 | 18.01 ± 0.62 | 0.02 ± 0.01 | 1.21 ± 0.03 | 16.90 ± 0.47 | 0.02 ± 0.00 | -1.85 | -2.09 | -0.3 | 0.032 (0.036*) | 0.006 (0.006*) | 0.821 (0.937) |
| **Cerebral WM** | 1.80 ± 0.09 | 21.28 ± 1.06 | 0.05 ± 0.01 | 1.65 ± 0.06 | 18.71 ± 0.75 | 0.05 ± 0.01 | -2.19 | -2.93 | -0.43 | <0.001 (<0.001*) | <0.001 (<0.001*) | 0.590 (0.937) |
| **Occipital–Parietal WM** | 1.75 ± 0.06 | 21.35 ± 1.51 | 0.06 ± 0.01 | 1.57 ± 0.07 | 17.61 ± 0.65 | 0.05 ± 0.01 | -2.83 | -3.52 | -0.44 | <0.001 (<0.001*) | <0.001 (<0.001*) | 0.462 (0.937) |
| **Temporo–Parietal WM** | 1.75 ± 0.06 | 21.06 ± 1.06 | 0.05 ± 0.01 | 1.58 ± 0.06 | 17.62 ± 0.70 | 0.05 ± 0.01 | -2.94 | -4.03 | -0.19 | <0.001 (<0.001*) | <0.001 (<0.001*) | 0.960 (0.960) |
| **Frontal WM** | 1.65 ± 0.10 | 20.46 ± 1.22 | 0.03 ± 0.01 | 1.58 ± 0.05 | 19.82 ± 0.99 | 0.03 ± 0.01 | -1.02 | -0.60 | -0.39 | 0.011 (0.014*) | 0.107 (0.114) | 0.398 (0.937) |
| **Cerebellar WM** | 1.60 ± 0.05 | 19.12 ± 0.90 | 0.06 ± 0.01 | 1.56 ± 0.07 | 17.89 ± 0.68 | 0.07 ± 0.01 | -0.58 | -1.60 | 0.41 | 0.200 (0.200) | 0.002 (0.002*) | 0.151 (0.937) |
| **CC** | 1.87 ± 0.09 | 21.28 ± 0.96 | 0.07 ± 0.01 | 1.77 ± 0.05 | 19.93 ± 0.79 | 0.07 ± 0.01 | -1.47 | -1.57 | -0.17 | <0.001 (<0.001*) | <0.001 (<0.001*) | 0.803 (0.937) |
| **AF** | 1.78 ± 0.11 | 21.87 ± 1.08 | 0.05 ± 0.01 | 1.62 ± 0.06 | 19.02 ± 0.88 | 0.05 ± 0.01 | -1.92 | -2.96 | -0.44 | <0.001 (<0.001*) | <0.001 (<0.001*) | 0.462 (0.937) |
| **CST** | 1.83 ± 0.08 | 20.69 ± 0.78 | 0.04 ± 0.01 | 1.67 ± 0.06 | 18.53 ± 1.15 | 0.03 ± 0.01 | -2.31 | -2.10 | -0.42 | <0.001 (<0.001*) | <0.001 (<0.001*) | 0.591 (0.937) |
| **IFOF** | 1.76 ± 0.08 | 21.88 ± 1.17 | 0.05 ± 0.01 | 1.63 ± 0.06 | 18.96 ± 0.59 | 0.05 ± 0.01 | -1.98 | -3.41 | -0.46 | <0.001 (<0.001*) | <0.001 (<0.001*) | 0.535 (0.937) |
| **ILF** | 1.73 ± 0.06 | 21.42 ± 1.09 | 0.06 ± 0.01 | 1.58 ± 0.06 | 17.88 ± 0.63 | 0.05 ± 0.01 | -2.43 | -4.25 | -0.30 | <0.001 (<0.001*) | <0.001 (<0.001*) | 0.761 (0.937) |
| **SLF1** | 1.74 ± 0.11 | 20.49 ± 1.14 | 0.05 ± 0.01 | 1.60 ± 0.04 | 18.48 ± 1.0 | 0.04 ± 0.01 | -1.81 | -1.91 | -0.21 | <0.001 (<0.001*) | <0.001 (<0.001*) | 0.757 (0.937) |
| **SLF2** | 1.78 ± 0.11 | 21.30 ± 1.14 | 0.05 ± 0.01 | 1.61 ± 0.05 | 18.81 ± 0.99 | 0.05 ± 0.01 | -2.20 | -2.38 | -0.45 | <0.001 (<0.001*) | <0.001 (<0.001*) | 0.379 (0.937) |
| **SLF3** | 1.76 ± 0.10 | 21.46 ± 1.04 | 0.05 ± 0.01 | 1.57 ± 0.06 | 18.57 ± 0.95 | 0.05 ± 0.01 | -2.52 | -2.92 | -0.2 | <0.001 (<0.001*) | <0.001 (<0.001*) | 0.826 (0.937) |

**Figure 6.**
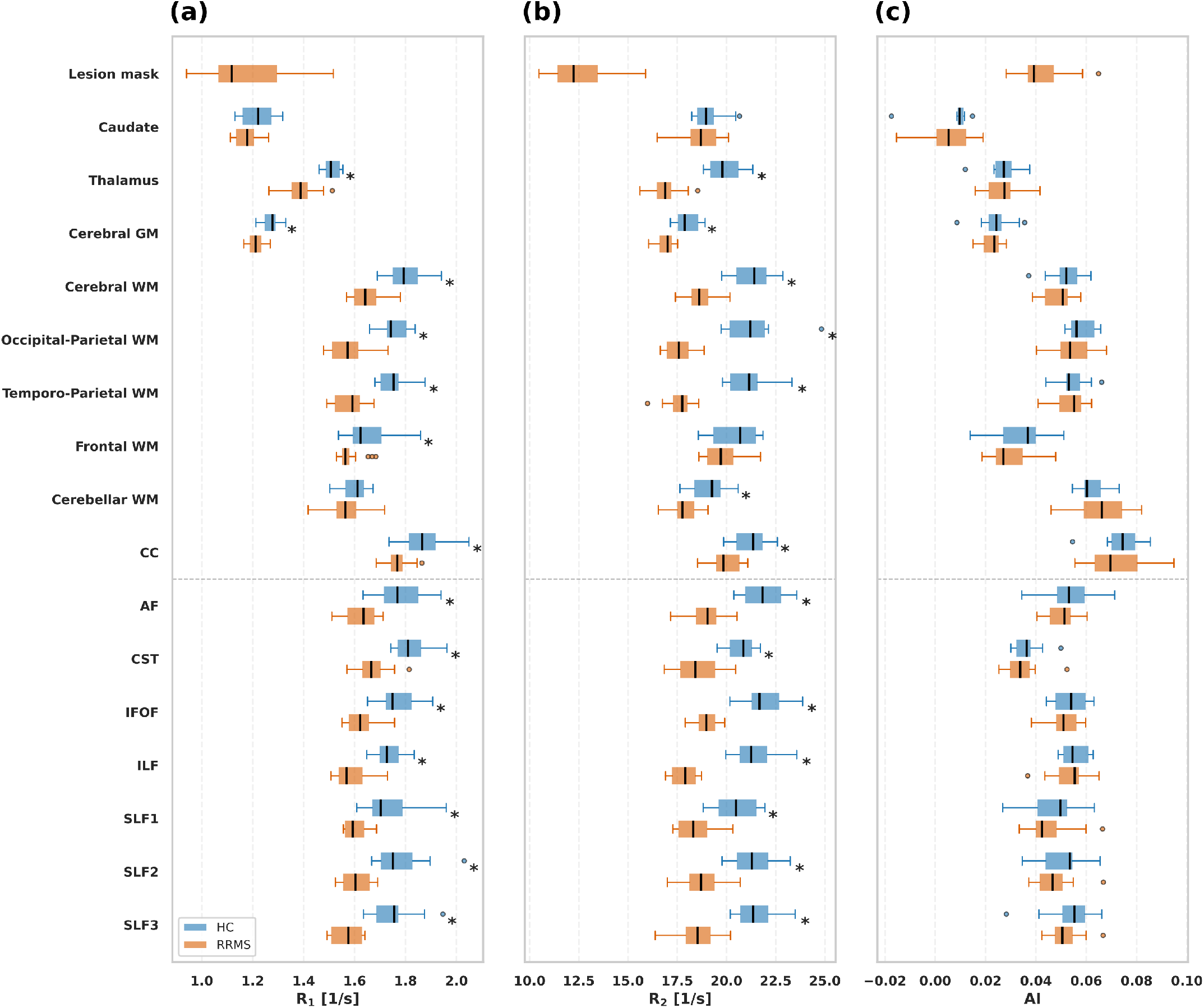
ROI analysis in NAWM. Boxplots for the mean *R*_1_ (a), *R*_2_ (b), and AI (c) values of NAWM in age-matched HC (blue) and RRMS (orange) cohorts are shown for all ROIs included in Figure 2, as well as for distinct WM bundles, i.e. superior longitudinal fasciculus (SLF1, SLF2, SLF3), inferior longitudinal fasciculus (ILF), inferior fronto-occipital fasciculus (IFOF), corticospinal tract (CST), arcuate fasciculus (AF), and the lesion mask. Asterisks (*) indicate ROIs with significant HC *−* RRMS differences after FDR correction (p_corr_ *<* 0.05).

## Discussion

This study demonstrates the first application of pc-bSSFP imaging in RRMS patients, revealing intermediate *R*_1_ and *R*_2_ relaxation rates in NAWM lying between the values observed in WM in HC and those observed in focal RRMS lesions. High intra- and inter-session repeatability in HC and detectable reduction of mean anisotropy in *R*_2_ and AI metrics for RRMS relative to HC were found.

### Repeatability and Anisotropy of pc-bSSFP Tissue Characterization

The biophysical origins of relaxation anisotropy are diverse, with potential contributors including the anisotropic magnetic susceptibility of myelin [40] or residual dipolar couplings within myelin sheaths [41]. *R*_2_ relaxation rates have been reported to increase with larger fiber-to-field angles *θ* [42] in agreement with our results. Since relaxation anisotropy reflects indirect myelin sensitivity, it can be used to differentiate age-related effects in WM structures [43] or pathological changes [44]. In this work, we investigate NAWM in RRMS patients as a use case for tissue anisotropy assessment, observing a slight global reduction in anisotropy, which may reflect subtle microstructural changes. The high scan-rescan repeatability of the AI supports the observed trend of reduced mean anisotropy in RRMS compared to HC. Comparable scan-rescan performance was achieved for *R*_1_ and *R*_2_ in this work as for another MP relaxometry framework [20].

### Normal Appearing White Matter and Tract Analysis

In agreement with previous studies reporting increased NAWM relaxation times (thus reduced relaxation rates) in RRMS patients [5, 11], this study demonstrates that NAWM *R*_1_ and *R*_2_ values in RRMS lie intermediate between focal lesions and HC WM relaxation rates. Increased *T*_1_ and *T*_2_ relaxation times have been associated with microstructural tissue alterations, including demyelination and chronic gliosis [8]. While this study did not explicitly relate total lesion burden to overall NAWM changes, mean differences between HC WM and NAWM in RRMS were significant for *R*_1_ and *R*_2_.

Microstructural characterization in MS patients has been shown to correlate more strongly with disability scores than conventional radiological biomarkers [45]. Further research should therefore aim at detailed WM tract-specific analyses to better delineate the impact of spatially localized microstructural changes on disability progression [45], as well as potential associations between quantitative MR metrics and underlying tis-sue damage [22, 23]. A recent study exploring high-resolution *T*_1_ mapping combined with WM tract analysis in a normative atlas space suggests that NAWM tissue alterations from qMRI may be more predictive of future disease activity than lesion load alone [45]. In the present work, z-score analysis in the MNI space was performed qualitatively, with similar whole-brain patterns observed in individual RRMS *R*_1_ and *R*_2_ maps compared to the results reported by Ravano et al. [45]. Moreover, this study demonstrates group differences between patients with RRMS and HC along WM tracts for *R*_1_ and *R*_2_, which was previously investigated primarily using DTI metrics or myelin water fraction [22, 23].

### Phase-Cycled bSSFP Imaging in RRMS

Other myelin-sensitive sequences, such as multiecho spin echo (MESE) [5], are optimized for three-compartment analysis and may improve differentiation of DAWM, NAWM, and focal lesions compared with the single-compartment approach used in this study. However, these sequences typically involve a trade-off in spatial resolution, resulting in relatively thick slices [5, 11, 46] (up to 8 mm [5]) to maintain clinically feasible scan times for the estimation of a single parameter.

Recent work has demonstrated MP frameworks for simultaneous *R*_1_ and *R*_2_ mapping in patients with MS [20, 21]. The significant findings reported by Ontaneda et al. [21] using MRF and successful disability progression prediction from thalamic volume in MS by Hänninen et al. [47] align with the large standardized effect sizes in the corpus callosum and thalamus as well as medium effects in the caudate, observed in the present study. Only medium effect sizes were observed for frontal NAWM *R*_2_ in this work. While pc-bSSFP imaging benefits from efficient 3D encoding and intrinsically high SNR, enabling high isotropic resolution with whole-brain coverage, transient-state MP frameworks typically require thicker slices ( ≈ 5 mm [20, 21]) to maintain sufficient SNR and achieve clinically feasible scan times. This limitation may reduce sensitivity to smaller lesions and subtle DAWM regions.

### Limitations

A major limitation of this study is the lack of covariate information, including longitudinal data, age stratification, and detailed clinical measures. Longitudinal studies combined with disability scores such as the expanded disability status scale (EDSS), may provide further insights into disease progression [46] and the transformation of DAWM into focal lesions in RRMS patients [48]. In addition, quantitative metrics such as *R*_2_ have been shown to be age-dependent [43], making the investigation of age-related effects important for future studies. Moreover, increasing the sample size will improve the explanatory power of the proposed analyses and could be facilitated by integrating the protocol into clinical routine. To this end, suitable 3D undersampling strategies and time-efficient alternatives like frequency-modulated SSFP [49] need to be evaluated in the future to accelerate the current ≈ 10 min pc-bSSFP protocol, which exhibited residual motion sensitivity leading to the exclusion of four RRMS subjects in this study. Another limitation relates to the use of single-shell DTI data for estimating crossing fibers and performing tractography. Multi-shell diffusion models may provide more robust fiber tracking analysis in particular in crossing fiber scenarios [50].

## Conclusion

Quantitative and qualitative analyses in this work suggest that pc-bSSFP-derived relaxometry metrics are sensitive to microstructural changes in NAWM. Moreover, the anisotropy of the assessed quantitative metrics was reproducible in the analyzed dataset, with a measurable reduction observed in NAWM of RRMS patients. Phase-cycled bSSFP provides an SNR-efficient approach for probing microstructural tissue properties, with potential for further acceleration strategies. In combination with fast, flexible, and robust fitting algorithms for MP mapping [27], this method may enable novel insights into the etiopathogenesis of neurodegenerative diseases such as MS.

## ACKNOWLEDGEMENTS

This research was supported by DFG Grant HE 9297/1-1. We gratefully acknowledge Silke Buschbach and Uta Groepers for their assistance with the MRI measurements of the RRMS participants.

For the purpose of open access, the authors have applied a Creative Commons Attribution (CC BY) license to any Author Accepted Manuscript version arising.

## References

[1] James Cairns et al. “Diffusely abnormal white matter in multiple sclerosis”. en. In: Journal of Neuroimaging 32.1 (Jan. 2022), pp. 5–16. ISSN: 1051-2284, 1552-6569. DOI: 10.1111/jon.12945.

[2] Gema Muñoz González et al. “A focus on the normal-appearing white and gray matter within the multiple sclerosis brain: a link to smoldering progression”. en. In: Acta Neuropathologica 150.1 (Aug. 2025), p. 16. ISSN: 1432-0533. DOI: 10.1007/s00401-025-02923-1.

[3] Janne West et al. “Normal Appearing and Diffusely Abnormal White Matter in Patients with Multiple Sclerosis Assessed with Quantitative MR”. en. In: PLoS ONE 9.4 (Apr. 2014). Ed. by Pablo Villoslada, e95161. ISSN: 1932-6203. DOI: 10.1371/journal.pone.0095161.

[4] R. Davis Holmes et al. “Nonlesional diffusely abnormal appearing white matter in clinically isolated syndrome: Prevalence, association with clinical and MRI features, and risk for conversion to multiple sclerosis”. en. In: Journal of Neuroimaging 31.5 (Sept. 2021), pp. 981–994. ISSN: 1051-2284, 1552-6569. DOI: 10.1111/jon.12900.

[5] Efrosini Papadaki et al. “T2 Relaxometry Evidence of Microstructural Changes in Diffusely Abnormal White Matter in Relapsing–Remitting Multiple Sclerosis and Clinically Isolated Syndrome: Impact on Visuomotor Performance”. en. In: Journal of Magnetic Resonance Imaging 54.4 (Oct. 2021), pp. 1077–1087. ISSN: 1053-1807, 1522-2586. DOI: 10.1002/jmri.27661.

[6] Carmen Tur et al. “Association of Early Progression Independent of Relapse Activity With Long-term Disability After a First Demyelinating Event in Multiple Sclerosis”. en. In: JAMA Neurology 80.2 (Feb. 2023), p. 151. ISSN: 2168-6149. DOI: 10.1001/jamaneurol.2022.4655.

[7] Antonio Scalfari et al. “Smouldering-Associated Worsening in Multiple Sclerosis: An International Consensus Statement on Definition, Biology, Clinical Implications, and Future Directions”. en. In: Annals of Neurology 96.5 (Nov. 2024), pp. 826–845. ISSN: 0364-5134, 1531-8249. DOI: 10.1002/ana.27034.

[8] Alexandra Seewann et al. “Diffusely Abnormal White Matter in Chronic Multiple Sclerosis: Imaging and Histopathologic Analysis”. en. In: Archives of Neurology 66.5 (May 2009). ISSN: 0003-9942. DOI: 10.1001/archneurol.2009.57.

[9] H. Vrenken, A. Seewann, D.L. Knol, C.H. Polman, F. Barkhof, and J.J.G. Geurts. “Diffusely Abnormal White Matter in Progressive Multiple Sclerosis: In Vivo Quantitative MR Imaging Characterization and Comparison between Disease Types”. en. In: American Journal of Neuroradiology 31.3 (Mar. 2010), pp. 541–548. ISSN: 0195-6108, 1936-959X. DOI: 10.3174/ajnr.A1839.

[10] Reza Rahmanzadeh et al. “Myelin and axon pathology in multiple sclerosis assessed by myelin water and multi-shell diffusion imaging”. en. In: Brain 144.6 (July 2021), pp. 1684–1696. ISSN: 0006-8950, 1460-2156. DOI: 10.1093/brain/awab088.

[11] James G. Harper et al. “Quantitative T1 brain mapping in early relapsing-remitting multiple sclerosis: longitudinal changes, lesion heterogeneity and disability”. en. In: European Radiology 34.6 (Nov. 2023), pp. 3826–3839. ISSN: 1432-1084. DOI: 10.1007/s00330-023-10351-6.

[12] Nikolaus Weiskopf et al. “Quantitative multi-parameter mapping of R1, PD*, MT, and R2* at 3T: a multi-center validation”. In: Frontiers in Neuroscience 7 (2013). ISSN: 1662-453X. DOI: 10.3389/fnins.2013.00095.

[13] Damien Nguyen and Oliver Bieri. “Motion-Insensitive Rapid Configuration Relaxometry”. en. In: Magnetic Resonance in Medicine 78.2 (Aug. 2017), pp. 518–526. ISSN: 07403194. DOI: 10.1002/mrm.26384.

[14] Yulia Shcherbakova, Cornelis A.T. van den Berg, Chrit T.W. Moonen, and Lambertus W. Bartels. “PLANET: An ellipse fitting approach for simultaneous T 1 and T 2 mapping using phase-cycled balanced steady-state free precession”. en. In: Magnetic Resonance in Medicine 79.2 (Feb. 2018), pp. 711–722. ISSN: 07403194. DOI: 10.1002/mrm.26717.

[15] Nils M. J. Plähn et al. “ORACLE: An analytical approach for T1, T2, proton density, and off-resonance mapping with phase-cycled balanced steady-state free precession”. en. In: Magnetic Resonance in Medicine (Dec. 2024), mrm.30388. ISSN: 0740-3194, 1522-2594. DOI: 10.1002/mrm.30388.

[16] Dan Ma et al. “Fast 3D magnetic resonance fingerprinting for a whole-brain coverage”. en. In: Magnetic Resonance in Medicine 79.4 (Apr. 2018), pp. 2190–2197. ISSN: 0740-3194, 1522-2594. DOI: 10.1002/mrm.26886.

[17] Tianle Cao et al. “Three-dimensional simultaneous brain mapping of T1, T2, and magnetic susceptibility with MR Multitasking”. en. In: Magnetic Resonance in Medicine 87.3 (Mar. 2022), pp. 1375–1389. ISSN: 0740-3194, 1522-2594. DOI: 10.1002/mrm.29059.

[18] Hongyan Liu et al. “A three-dimensional Magnetic Resonance Spin Tomography in Time-domain protocol for high-resolution multiparametric quantitative magnetic resonance imaging”. en. In: NMR in Biomedicine 37.2 (Feb. 2024), e5050. ISSN: 0952-3480, 1099-1492. DOI: 10.1002/nbm.5050.

[19] Nora Vandeleene et al. “Using quantitative magnetic resonance imaging to track cerebral alterations in multiple sclerosis brain: A longitudinal study”. en. In: Brain and Behavior 13.5 (May 2023), e2923. ISSN: 2162-3279, 2162-3279. DOI: 10.1002/brb3.2923.

[20] Sen Ma et al. “Three-dimensional simultaneous brain T1, T2, and ADC mapping with MR Multitasking”. en. In: Magnetic Resonance in Medicine 84.1 (July 2020), pp. 72–88. ISSN: 0740-3194, 1522-2594. DOI: 10.1002/mrm.28092.

[21] Daniel Ontaneda et al. “Magnetic resonance fingerprinting in multiple sclerosis”. en. In: Multiple Sclerosis and Related Disorders 79 (Nov. 2023), p. 105024. ISSN: 22110348. DOI: 10.1016/j.msard.2023.105024.

[22] Michael Dayan et al. “Profilometry: A new statistical framework for the characterization of white matter pathways, with application to multiple sclerosis”. en. In: Human Brain Mapping 37.3 (Mar. 2016), pp. 989–1004. ISSN: 1065-9471, 1097-0193. DOI:10.1002/hbm.23082.

[23] Erick Hernandez-Gutierrez et al. “Multi-tensor fixel-based metrics in tractometry: application to multiple sclerosis”. In: Frontiers in Neuroscience 18 (Dec. 2024), p. 1467786. ISSN: 1662-453X. DOI: 10.3389/fnins.2024.1467786.

[24] Klaus Scheffler and Stefan Lehnhardt. “Principles and applications of balanced SSFP techniques”. en. In: European Radiology 13.11 (Nov. 2003), pp. 2409–2418. ISSN: 0938-7994, 1432-1084.

[25] Karla L. Miller, Stephen M. Smith, and Peter Jezzard. “Asymmetries of the balanced SSFP profile. Part II: White matter”. In: Magnetic Resonance in Medicine 63.2 (Feb. 2010), pp. 396–406. DOI: 10.1002/mrm.22249.

[26] Rahel Heule, Jonas Bause, Orso Pusterla, and Klaus Scheffler. “Multi-parametric artificial neural network fitting of phase-cycled balanced steady-state free precession data”. en. In: Magnetic Resonance in Medicine 84.6 (Dec. 2020), pp. 2981–2993. ISSN: 0740-3194, 1522-2594. DOI: 10.1002/mrm.28325.

[27] Florian Birk, Lucas Mahler, Julius Steiglechner, Qi Wang, Klaus Scheffler, and Rahel Heule. “Flexible and cost-effective deep learning for accelerated multi-parametric relaxometry using phase-cycled bSSFP”. en. In: Scientific Reports 15.1 (Feb. 2025), p. 4825. ISSN: 2045-2322. DOI: 10.1038/s41598-025-88579-z.

[28] Giulia M. C. Rossi et al. “SPARCQ: A new approach for fat fraction mapping using asymmetries in the phase-cycled balanced SSFP signal profile”. en. In: Magnetic Resonance in Medicine 90.6 (Dec. 2023), pp. 2348–2361. ISSN: 0740-3194, 1522-2594. DOI: 10.1002/mrm.29813.

[29] Florian Birk et al. “High-resolution neural network-driven mapping of multiple diffusion metrics leveraging asymmetries in the balanced steady-state free precession frequency profile”. en. In: NMR in Biomedicine (Dec. 2021), e4669. ISSN: 0952-3480, 1099-149. DOI: 10.1002/nbm.4669.

[30] John P. Mugler and James R. Brookeman. “Three-dimensional magnetization-prepared rapid gradient-echo imaging (3D MP RAGE)”. en. In: Magnetic Resonance in Medicine 15.1 (July 1990), pp. 152–157. ISSN: 0740-3194, 1522-2594. DOI: 10.1002/mrm.1910150117.

[31] Joseph V. Hajnal et al. “Use of Fluid Attenuated Inversion Recovery (FLAIR) Pulse Sequences in MRI of the Brain:” en. In: Journal of Computer Assisted Tomography 16.6 (Nov. 1992), pp. 841–844. ISSN: 0363-8715. DOI: 10.1097/00004728-199211000-00001.

[32] Sohae Chung, Daniel Kim, Elodie Breton, and Leon Axel. “Rapid B1 + mapping using a preconditioning RF pulse with TurboFLASH readout”. en. In: Magnetic Resonance in Medicine 64.2 (Aug. 2010), pp. 439–446. ISSN: 0740-3194, 1522-2594. DOI: 10.1002/mrm.22423.

[33] Tobias Lindig et al. “Proof of principle for the clinical use of a CE-certified automatic imaging analysis tool in rare diseases studying hereditary spastic paraplegia type 4 (SPG4)”. en. In: Scientific Reports 12.1 (Dec. 2022), p. 22075. ISSN: 2045-2322. DOI:10.1038/s41598-022-25545-z.

[34] Guillaume Theaud, Jean-Christophe Houde, Arnaud Boré, François Rheault, Felix Morency, and Maxime Descoteaux. “TractoFlow: A robust, efficient and reproducible diffusion MRI pipeline leveraging Nextflow & Singularity”. en. In: NeuroImage 218 (Sept. 2020), p. 116889. ISSN: 10538119. DOI: 10.1016/j.neuroimage.2020.116889.

[35] Etienne St-Onge, Kurt G Schilling, and Francois Rheault. BundleSeg: A versatile, reliable and reproducible approach to white matter bundle segmentation. Version Number: 1. 2023. DOI: 10.48550/ARXIV.2308.10958.

[36] Martin Cousineau et al. “A test-retest study on Parkinson’s PPMI dataset yields statistically significant white matter fascicles”. en. In: NeuroImage: Clinical 16 (2017), pp. 222–233. ISSN: 22131582. DOI: 10.1016/j.nicl.2017.07.020.

[37] Fang-Cheng Yeh. “Population-based tract-to-region connectome of the human brain and its hierarchical topology”. en. In: Nature Communications 13.1 (Aug. 2022), p. 4933. ISSN: 2041-1723. DOI: 10.1038/s41467-022-32595-4.

[38] Elias Kellner, Bibek Dhital, Valerij G. Kiselev, and Marco Reisert. “Gibbs-ringing artifact removal based on local subvoxel-shifts: Gibbs-Ringing Artifact Removal”. en. In: Magnetic Resonance in Medicine 76.5 (Nov. 2016), pp. 1574–1581. ISSN: 07403194. DOI: 10.1002/mrm.26054.

[39] Malte Hoffmann, Benjamin Billot, Douglas N. Greve, Juan Eugenio Iglesias, Bruce Fischl, and Adrian V. Dalca. “SynthMorph: learning contrast-invariant registration without acquired images”. In: IEEE Transactions on Medical Imaging 41.3 (Mar. 2022). arXiv:2004.10282 [eess], pp. 543–558. ISSN: 0278-0062, 1558-254X. DOI: 10.1109/TMI.2021.3116879.

[40] Samuel Wharton and Richard Bowtell. “Gradient echo based fiber orientation mapping using R2* and frequency difference measurements”. en. In: NeuroImage 83 (Dec. 2013), pp. 1011–1023. ISSN: 10538119. DOI: 10.1016/j.neuroimage.2013.07.054.

[41] Yuxi Pang. “Orientation dependent proton transverse relaxation in human brain white matter: The magic angle effect on a cylindrical helix”. en. In: Magnetic Resonance Imaging 100 (July 2023), pp. 73–83. ISSN: 0730725X. DOI: 10.1016/j.mri.2023.03.010.

[42] Chantal M.W. Tax, Elena Kleban, Maxime Chamberland, Muhamed Baraković, Umesh Rudrapatna, and Derek K. Jones. “Measuring compartmental T 2 - orientational dependence in human brain white matter using a tiltable RF coil and diffusion-T 2 correlation MRI”. en. In: NeuroImage 236 (Aug. 2021), p. 117967. ISSN: 10538119. DOI: 10.1016/j.neuroimage.2021.117967.

[43] Michael J. Knight, Serena Dillon, Lina Jarutyte, and Risto A. Kauppinen. “Magnetic Resonance Relaxation Anisotropy: Physical Principles and Uses in Microstructure Imaging”. en. In: Biophysical Journal 112.7 (Apr. 2017), pp. 1517–1528. ISSN: 00063495. DOI: 10.1016/j.bpj.2017.02.026.

[44] Enedino Hernández-Torres et al. “Orientation Dependent MR Signal Decay Differentiates between People with MS, Their Asymptomatic Siblings and Unrelated Healthy Controls”. en. In: PLOS ONE 10.10 (Oct. 2015). Ed. by Orhan Aktas, e0140956. ISSN: 1932-6203. DOI: 10.1371/journal.pone.0140956.

[45] Veronica Ravano et al. “Tract-wise microstructural analysis informs on current and future disability in early multiple sclerosis”. en. In: Journal of Neurology 271.2 (Feb. 2024), pp. 631–641. ISSN: 0340-5354, 1432-1459. DOI: 10.1007/s00415-023-12023-3.

[46] Pietro Bontempi et al. “Non-lesional white matter in relapsing–remitting multiple sclerosis assessed by multicomponent T2 relaxation”. en. In: Brain and Behavior 13.12 (Dec. 2023), e3334. ISSN: 2162-3279, 2162-3279. DOI: 10.1002/brb3.3334.

[47] Katariina Hänninen et al. “Thalamic Atrophy Predicts 5-Year Disability Progression in Multiple Sclerosis”. In: Frontiers in Neurology 11 (July 2020), p. 606. ISSN: 1664-2295. DOI: 10.3389/fneur.2020.00606.

[48] Mahsa Dadar, Sawsan Mahmoud, Sridar Narayanan, D Louis Collins, Douglas L Arnold, and Josefina Maranzano. “Diffusely abnormal white matter converts to T2 lesion volume in the absence of MRI-detectable acute inflammation”. en. In: Brain 145.6 (June 2022), pp. 2008–2017. ISSN: 0006-8950, 1460-2156. DOI: 10.1093/brain/awab448.

[49] Volkert Roeloffs, Sebastian Rosenzweig, H. Christian M. Holme, Martin Uecker, and Jens Frahm. “Frequency-modulated SSFP with radial sampling and subspace reconstruction: A time-efficient alternative to phase-cycled bSSFP”. en. In: Magnetic Resonance in Medicine 81.3 (Mar. 2019), pp. 1566–1579. ISSN: 0740-3194, 1522-2594. DOI: 10.1002/mrm.27505.

[50] Colin Vanden Bulcke, Anna Stölting, Dragan Maric, Benoît Macq, Martina Absinta, and Pietro Maggi. “Comparative overview of multi-shell diffusion MRI models to characterize the microstructure of multiple sclerosis lesions and periplaques”. en. In: NeuroImage: Clinical 42 (2024), p. 103593. ISSN: 22131582. DOI: 10.1016/j.nicl.2024.103593.

